# Genome-wide development of lncRNA-derived-SSR markers for Dongxiang wild rice (*Oryza rufipogon* Griff.)

**DOI:** 10.1101/2021.08.23.457289

**Authors:** Wanling Yang, Yuanwei Fan, Yong Chen, Gumu Ding, Hu Liu, Jiankun Xie, Fantao Zhang

## Abstract

Dongxiang wild rice (*Oryza rufipogon* Griff., DXWR) is the northernmost distributed common wild rice found in the world. It contains a large number of agronomically valuable genes, which makes it a natural gene pool for rice breeding. Molecular markers, especially simple repeat sequence (SSR) markers, play important roles in crop breeding. Although a large number of SSR markers have been developed, most of them are derived from the genome coding sequences, rarely from non-coding sequences. Meanwhile, long non-coding RNAs (lncRNAs), which are derived from the transcription of non-coding sequences, play vital roles in plant growth, development and stress responses. In this study, 1878 SSR loci were detected from the lncRNA sequences of DXWR, and 1258 lncRNA-derived-SSR markers were developed on the genome-wide scale. To verify the validity and applicability of these markers, 72 pairs of primers were randomly selected to test 44 rice materials. The results showed that 42 (58.33%) pairs of primers have abundant polymorphism among these rice materials; the polymorphism information content (PIC) values ranged from 0.04 to 0.87 with an average of 0.50; the genetic diversity index of SSR loci varied from 0.04 to 0.88 with an average of 0.56; and the number of alleles per marker ranged from 2 to 11 with an average of 4.36. Thus, we concluded that these lncRNA-derived-SSR markers are a very useful source for future basic and applied research, including genetic diversity analysis, QTL mapping, and molecular breeding programs, to make good use of the elite lncRNA genes from DXWR.

## Introduction

Dongxiang wild rice (*Oryza rufipogon* Griff., DXWR) is a common wild rice species found in Dongxiang County, Jiangxi Province, China. It is by far the northernmost (28°14’ N) distributed common wild rice found in the world (Mao *et al.*, 2015). Previous studies have showed that DXWR possesses many stress tolerance and high yield-related genes, which have been lost in modern cultivated rice (Zhang *et al.*, 2016). Therefore, DXWR is an elite genetic resource to breed new rice varieties.

In eukaryotes, over 90% of the genome is transcribed to large amounts of non-coding RNAs (ncRNAs) that are not translated into proteins (Wierzbicki, 2012). Long non-coding RNAs (lncRNAs) are an important type of ncRNAs. In plants, lncRNAs act as key regulators of plant growth and development, as well as in response to various biotic and abiotic stresses (Xu *et al.*, 2017). Molecular marker-assisted selection (MAS) can make great use of the valuable genetic resources in wild rice by precisely selecting genes for various traits, therefore producing superior germplasm (Collard and Mackill, 2008; Akhtamov *et al.*, 2020). Simple repeat sequences (SSRs), also known as microsatellites, are DNA segments with 1-6 nucleotides short base-pair motif repeated several times in tandem (Kozlowski *et al.*, 2010). They have been extensively used in genetic diversity analysis and MAS breeding because of their high reproducibility, abundant polymorphisms, co-dominant inheritance, high genome coverage and simple analysis methods (Xie *et al.*, 2017; Singh *et al.*, 2018). Although recently we developed SSR markers from drought-stress responsive miRNA and identified seed storability-related loci in DXWR using whole-genome sequencing (Zhao *et al.*, 2021; Chen *et al.*, 2021), there are no reports on the development of lncRNA-derived-molecular markers in common wild rice so far, which greatly limits the discovery and utilization of the elite lncRNA genes from the important and endangered germplasm resource. Therefore, the main objectives of this study were to: (i) use the lncRNAs sequencing data to develop a set of lncRNA-derived-SSR markers and (ii) test their stabilities and polymorphisms for DXWR.

## Experimental

The information of rice materials used in this study was presented in Supplementary Table S1. To develop more lncRNA-derived-SSR markers, SSR Hunter software was used to identify all possible di-, tri-, tetra-, penta-, and hexa-nucleotide SSRs with a minimum set of three repeats, respectively (Li and Wan, 2005). Subsequently, primers were designed based on the flanking sequences of the SSRs using Primer3.0 software (Untergasser *et al.*, 2012). The main parameters of the primer design were as follows: the primer length was 18-25 bp, GC content was 40-60 %, melting temperature was 55-65 °C, the expected length of the amplification product was 100-250 bp.

Genomic DNA was extracted using the Plant Genomic DNA Rapid Extraction Kit (Sangon Biotech Co., Ltd). The PCR reaction system was 10 μL, including 1 μL (200 ng/μL) genomic DNA, 5 μL 2 × FastTaq Premix (Tolo Biotech Co., Ltd), 1 μL (0.01 nmol/μL) primers and 3 μL ddH2O. The PCR amplification reaction program was as follows: pre-denaturation at 95°C for 5 min; denaturation at 95°C for 30 s, annealing at 55°C for 45 s, extension at 72°C for 45 s for 30 cycles; and a final extension at 72°C for 10 min. The PCR products were run on 9 % denaturing polyacrylamide gel with 0.5 × TBE buffer. After electrophoresis, the gels were visualized using the silver staining method (Cook *et al.*, 2004). The genetic parameters were analyzed using Powermarker software (Liu and Muse, 2005).

## Discussion

In our previous study, a total of 1655 lncRNA transcripts were obtained from DXWR using strand-specific RNA sequencing (Qi *et al.*, 2020). In this study, to develop a set of lncRNA-derived-SSR markers for DXWR, SSR Hunter software was used to search for SSRs present in these lncRNA transcripts. Totally, 1878 SSRs were detected. Among them, the dinucleotide (1291, 68.74%) and trinucleotide (498, 26.52%) repeat motifs were the most abundant types (Table 1). Moreover, the types of AG/CT (505, 39.12%) and CCG/CGG (77, 15.46%) were the most predominant in the dinucleotide and trinucleotide repeat types, respectively (Fig. 1). These results following the same pattern as the lncRNA-derived-SSR study in wheat (Bhandawat *et al.*, 2020).

**Fig. 1.**
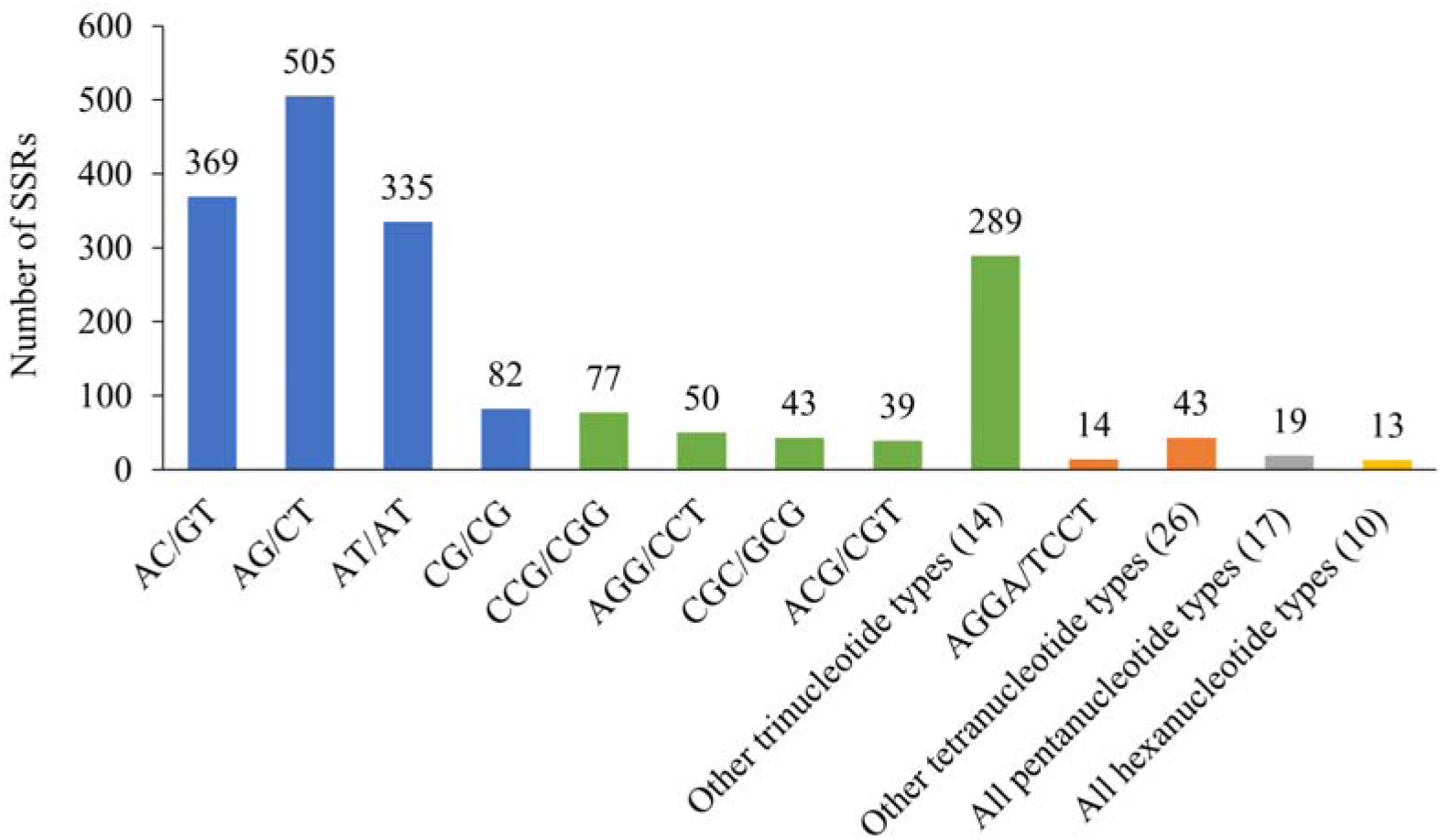
Frequency of different SSR repeat motif types in DXWR.

**Table 1.**
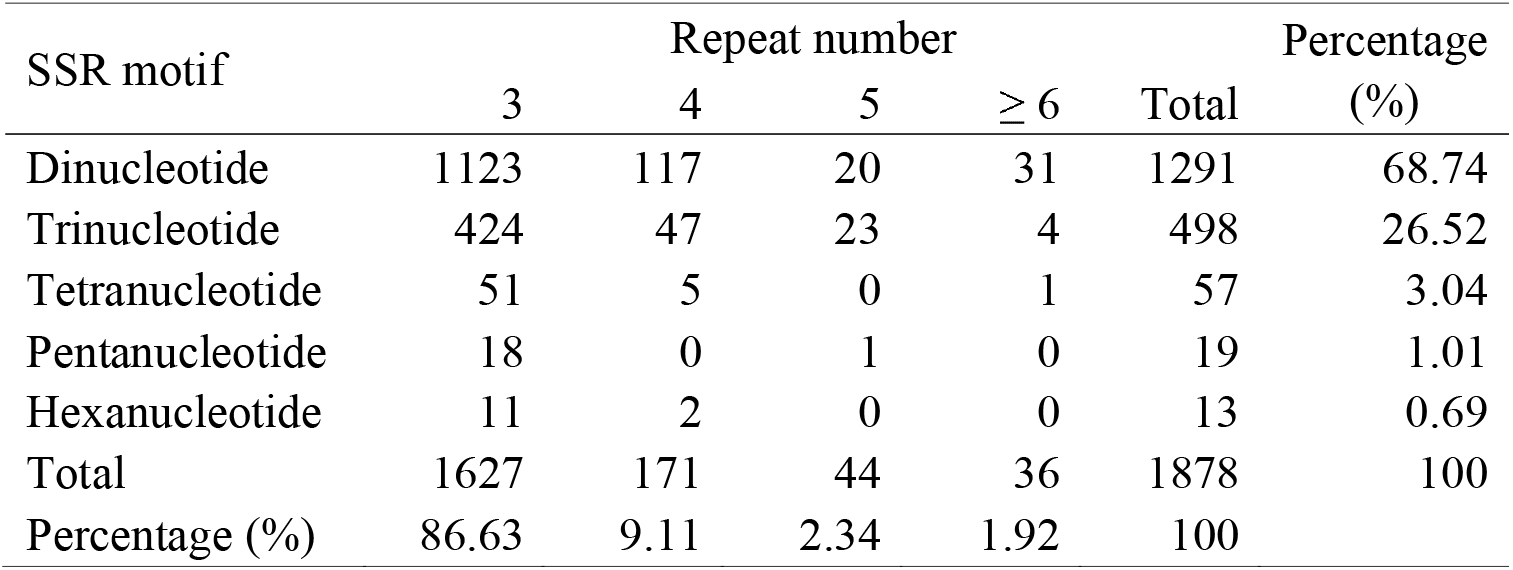
Percentage of the identified SSR motifs in different repeat types of DXWR

Based on the detected lncRNA-derived-SSR loci, we successfully developed a total of 1258 molecular markers, including 885 (70.35%) for dinucleotide repeat type, 316 (25.12%) for trinucleotide repeat type, 35 (2.78%) for tetranucleotide repeat type, 16 (1.27%) for pentanucleotide repeat type and 6 (0.48%) for hexanucleotide repeat type (Supplementary Table S2). These lncRNA-derived-SSR markers were present throughout all of the 12 chromosomes. Of these markers, 781 (62.08%) were found to be present in the first six chromosomes (Chr. 1-6); while 477 (37.92%) were present in the remaining six chromosomes (Chr. 7-12). This is largely due to the identified SSR loci exhibit some preference for the lncRNA transcripts located in chromosomes 1-6 of DXWR. It was also observed that Chr. 2 has the highest number of lncRNA-derived-SSR markers (169, 13.43%) while Chr.9 has the lowest number of lncRNA-derived-SSR markers (66, 5.25%).

To test the stabilities and polymorphisms of the developed lncRNA-derived-SSR markers, we randomly selected 72 pairs of primers for further analysis. As a result, 42 (58.33%) pairs of primers showed abundant polymorphisms among the 44 rice materials (Fig. 2). The 42 polymorphic SSRs produced a total of 183 alleles among the 44 rice materials, ranging from 2 to 11, with an average 4.36 alleles per locus. The polymorphic information content (PIC) of these polymorphic SSR markers ranged from 0.04 to 0.87 with an average of 0.50. The genetic diversity index was observed from 0.04 to 0.88, with a mean value of 0.56 (online Supplementary Table S3). The mean number of alleles and PIC value of the verified lncRNA-derived-SSR markers in our study were more than capsicum which were 2.50 and 0.39 respectively (Jaiswal *et al.*, 2020), indicating that the developed lncRNA-derived-SSR markers were highly polymorphic and can be wildly applied in DXWR and modern cultivated rice.

**Fig. 2.**
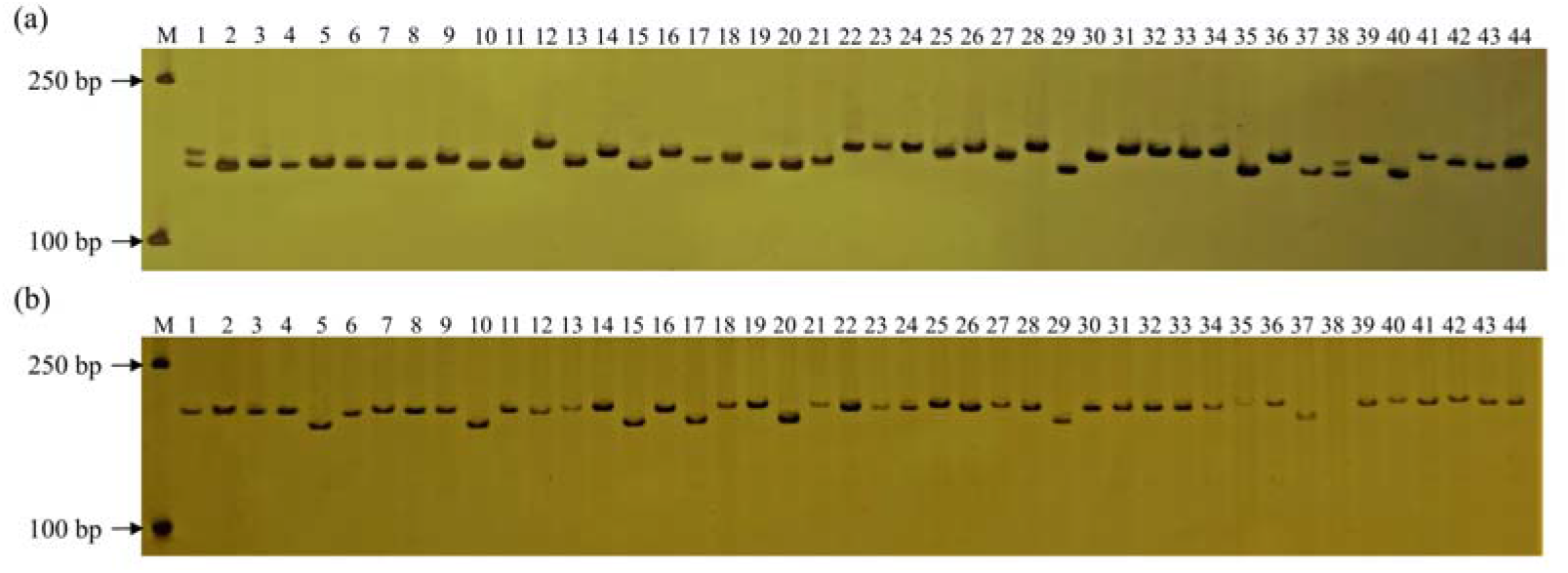
Representative results of polymorphic lncRNA-derived-SSR markers amplified by the genome of DXWR and other worldwide rice cultivars. (a) Lnc-SSR-566686; (b): Lnc-SSR-353993; M: DL2000 DNA Marker; 1-3: three different populations of DXWR; 4-44: other worldwide cultivars. The information of the 44 rice materials was presented in online Supplementary Table S1.

In summary, this is the first report of the development and characterization of lncRNA-derived-markers in common wild rice, which lays the foundation for discovery and utilization of the elite lncRNA genes to further make good use of the valuable and endangered wild rice germplasm resource.

## Acknowledgements

This research was partially supported by the National Natural Science Foundation of China (32070374, 31960370, 31960085), the Natural Science Foundation of Jiangxi Province, China (20202ACB205002), the Foundation of Jiangxi Provincial Key Lab of Protection and Utilization of Subtropical Plant Resources (YRD201903).

---

**Table S1.** Dongxiang wild rice and worldwide rice cultivars used for verification of the developed lncRNA-derived-SSR markers in this study

**Table S2.** The information of developed lncRNA-derived-SSR markers for Dongxiang wild rice

**Table S3.** The verified lncRNA-derived-SSR markers and their amplification results with 44 rice materials in this study

## Notes

### Competing Interest Statement

The authors have declared no competing interest.

